# A systematic survey of an intragenic epistatic landscape

**DOI:** 10.1101/010645

**Authors:** Claudia Bank, Ryan T. Hietpas, Jeffrey D. Jensen, Daniel N.A. Bolon

## Abstract

Mutations are the source of evolutionary variation. The interactions of multiple mutations can have important effects on fitness and evolutionary trajectories. We have recently described the distribution of fitness effects of all single mutations for a nine amino acid region of yeast Hsp90 (Hsp82) implicated in substrate binding. Here, we report and discuss the distribution of intragenic epistatic effects within this region in seven Hsp90 point mutant backgrounds of neutral to slightly deleterious effect, resulting in an analysis of more than 1000 double-mutants. We find negative epistasis between substitutions to be common, and positive epistasis to be rare – resulting in a pattern that indicates a drastic change in the distribution of fitness effects one step away from the wild type. This can be well explained by a concave relationship between phenotype and genotype (i.e., a concave shape of the local fitness landscape), suggesting mutational robustness intrinsic to the local sequence space. Structural analyses indicate that, in this region, epistatic effects are most pronounced when a solvent-inaccessible position is involved in the interaction. In contrast, all 18 observations of positive epistasis involved at least one mutation at a solvent-exposed position. By combining the analysis of evolutionary and biophysical properties of an epistatic landscape, these results contribute to a more detailed understanding of the complexity of protein evolution.

## INTRODUCTION

Mutation is the source of evolutionary variation, and over immense timescales, the cumulative effects of mutations have given rise to an enormous diversity of life. New mutations may be grouped into three general categories based upon their effect on organismal fitness: deleterious, neutral, and beneficial. Previous experimental studies indicate that the majority of new mutations in a standard environment are nearly neutral or strongly deleterious, with only a small minority conferring a fitness benefit (e.g., Sanjuán et al. 2004; Hietpas et al. 2011; Hietpas et al. 2013), consistent with population genetic theory (Kimura 1968). This results in a distribution of fitness effects (DFE) that is bimodal, with one peak centered at wild-type-like fitness, and another at (near-) lethality.

However, this work has been largely focused on the effects of single nucleotide changes. Whereas the simultaneous occurrence of two or more new mutations within a single gene may be rare in populations of moderate effective size and mutation rate, over many replicative events multiple mutations can accumulate on the same genetic background with potentially important consequences on, for example, the individual probabilities of fixation (Hill and Robertson 1966; Campos 2004; de Oliveira et al. 2008; Kermany and Lessard 2012). This was first considered by Bateson (Bateson 1909), who coined the term epistasis in what was one of the first joint considerations of Darwinian evolution with Mendelian genetics. Indeed, while such epistatic effects may be central to evolution, over 100 years later their study remains challenging to investigate in most experimental systems, owing to combinatorial complexity (Lehner 2011).

Yet, recent advances in sequencing technology enable us for the first time to study thousands of mutational interactions in a systematic and accurate way, thereby affording a direct approach to assess the scale of epistasis. Specifically, mutations may be considered independent if the fitness (or, more accurately, a phenotype that is measured as a proxy for fitness) of the combined mutant equals the product of the fitness of each individual mutant. Combinations of mutations that deviate from this rule are considered interdependent or epistatic. The interdependent fitness effects of epistatic mutations are directional and can result in combined mutants with fitness that is increased (referred to as *positive epistasis*) or decreased (*negative epistasis*) relative to independence.

Compensatory mutations - a form of positive epistasis in which one single mutation is deleterious, but a second, despite being neutral or deleterious on its own, brings an individual back to wild-type fitness - represent a form of epistasis with biologically and medically important ramifications. Many studies have demonstrated that compensatory mutations play an important role in the adaptation of microbes and viruses to antibiotic or antiviral treatment (e.g., Bloom et al. 2010; Abed et al. 2011; Martínez et al. 2011; Das et al. 2013). In many of these cases, the primary drug resistance mutation was found to be deleterious in the absence of drug, but a secondary mutation with a neutral fitness effect in the parental background increased the fitness of the primary mutation, promoting the persistence of the resistance mutation in the absence of drug treatment. A recent meta-analysis indicates that 83% of all compensatory mutations occur within the same gene as the primary mutation, emphasizing the relevance of intragenic epistasis in evolution (Poon et al. 2005).

Intragenic epistasis itself has a rich history of investigation in the framework of protein structure-function relationships. Double mutant cycles compare the biochemical properties of combined mutants to individual mutants and have proven a powerful approach to investigate the interdependence of mutations on protein stability or activity (Carter et al. 1984). These studies have been utilized to investigate a variety of intra-protein interactions including functional residues (Wall-Lacelle et al. 2011), long-range structural interactions (Istomin et al. 2008), exposed and buried salt bridges (Vaughan et al. 2002; Luisi et al. 2003), hydrogen bond networks (Jang et al. 2004), and protein-protein interactions (Schreiber and Fersht 1995). While double mutant cycles provide valuable biophysical and biochemical insights (Horovitz 1996), measuring the biochemical properties of many protein variants can be laborious, and the connections between biochemical function and fitness complex (Drummond et al. 2005; Gout et al. 2010; Jiang et al. 2013). Recently, a number of studies in experimentally evolved populations have indicated that the assessment of thermodynamic stability provides a key to identifying epistatic interactions, with a stabilizing mutation enabling a secondary mutation that improves protein function (Gong et al. 2013; Jacquier et al. 2013). Using random mutagenesis, an assessment of more than 100,000 mutations (and 40,000 double mutants) in an RNA recognition motif in *S. cerevisiae* yielded the largest and most systematic picture of intragenic epistasis to date (Melamed et al. 2013).

Epistasis has also been studied *in vivo*. For example, in yeast, the effects of specific mutations on fitness can be rapidly analyzed in the background of thousands of individual gene knockouts using the epistatic mini-array profile (E-MAP) approach (Schuldiner et al. 2005), or synthetic genetic analysis (SGA, Tong et al. 2001; van Opijnen et al. 2009). While epistasis mapping by these approaches has been extremely useful for detecting physiological connections between gene products, it is not well suited to investigate intragenic epistasis, or to comprehensively screen point mutants.

We previously developed an approach termed EMPIRIC to quantify the fitness effects of all possible point mutations in a gene or region of a gene (Hietpas et al. 2011; Hietpas et al. 2012; Roscoe et al. 2013) and used this approach to comprehensively delineate the distribution of fitness effects for a nine amino acid region of yeast Hsp90 (also known as Hsp82). Hsp90 is a homodimeric protein chaperone that plays an essential role in stress responses, kinase activation, and hormone receptor maturation (Nemoto et al. 1995; Pratt and Toft 2003; Zhao et al. 2005). To successfully perform these functions, Hsp90 binds to numerous co-chaperones during the process of substrate maturation. By 2-hybrid and SGA analysis, Hsp90 has been found to interact with ∼3% of the yeast proteome (Millson et al. 2004; Zhao et al. 2005).

The region of Hsp90 that is the focus of this work (amino acids 582-590) has many hallmarks of a putative substrate-binding interface (Harris et al. 2004; Ali et al. 2006; Hietpas et al. 2011; Street et al. 2014), including two positions with solvent exposed aromatic residues (F583 and W585). In our previous work, we observed that yeast fitness requires large hydrophobic amino acids at both of these positions (Hietpas et al. 2011; Jiang et al. 2013), indicating that they provide a critical hydrophobic docking site. We have also observed that a buried intra-molecular hydrogen bond mediated by S586 is critical for yeast fitness indicating that the main-chain conformation of this region is important for function. Together, these observations indicate that the physical properties of surface accessible side chains in this region of Hsp90 combined with main-chain conformational preferences directly impact fitness. To probe relationships between protein structure and epistasis in this region of Hsp90 we here utilize EMPIRIC to systematically determine fitness effects of point mutants within this region, on the backgrounds of seven local anchor mutations (Figure 1 and Table 1).

**Figure 1:**
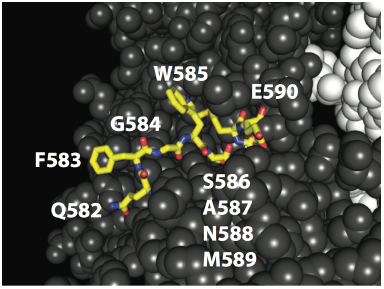
Structure of yeast Hsp90 illustrating the region investigated (amino acids 582-590) in yellow, containing wild-type amino acids with varied physical properties and buried and exposed positions as detailed in Table 1.

**Table 1.** (A) Positions studied and their biophysical properties

| Position | wild-type amino acid | % solvent accessible | Classification |
| --- | --- | --- | --- |
| 582 | Q | 54.69 | S<sup>a</sup> |
| 583 | F | 85.2 | S |
| 584 | G | 33.48 | S |
| 585 | W | 64.5 | S |
| 586 | S | 2.94 | C |
| 587 | A | 3.82 | C |
| 588 | N | 1.64 | C |
| 589 | M | 10.43 | C |
| 590 | E | 29.79 | S |

(B) Anchor mutations, their single-mutation growth rates and 95% credibility intervals
| Position | Amino acid | Median growth rate | 2.5% CrI | 97.5% CrI |
| --- | --- | --- | --- | --- |
| 583 | N | 0.9867 | 0.9797 | 0.9937 |
| 584 | F | 0.9761 | 0.9660 | 0.9858 |
| 584 | S | 0.9828 | 0.9764 | 0.9889 |
| 585 | L | 0.9813 | 0.9757 | 0.9871 |
| 586 | G | 0.9794 | 0.9742 | 0.9849 |
| 587 | G | 0.9913 | 0.9879 | 0.9947 |
| 588 | F | 1.0053 | 0.9988 | 1.0119 |

<sup>a</sup>Surface
<sup>b</sup>Core
<sup>c</sup>Credibility interval

Importantly, with our choice of wild-type-like to slightly deleterious anchor mutations, we cover a very different part of the sequence space than has been explored in previous systematic assessments of epistatic effects (e.g. Jacquier et al. 2013). Instead of anchoring a step towards a better-adapted phenotype (i.e., the “upwards” slope of the fitness landscape), we study the “downwards” fitness landscape - one step away from the optimal phenotype. This part of the fitness landscape becomes particularly important when considering small or bottlenecking populations in which drift may have a strong effect, such that slightly deleterious mutations have the potential to become fixed. Further, quantifying the changing shape of the DFE after a single mutational step allows for deeper insight in to both predictions of Fisher’s Geometric model (see Bank et al. 2014), as well as for notions of ‘mutational robustness’ (e.g., Draghi et al. 2010; Goldstein 2013; Lauring et al. 2013).

## RESULTS AND DISCUSSION

### Characterization of epistatic effects

We examined the general pattern of epistasis by comparing the expected growth rates under the assumption of multiplicative mutational effects with the observed growth rates of double mutants (cf. Fig. 2). Despite the conservative cutoff (see Materials and Methods), we observe ubiquitous and strong negative epistasis. Among all 1,015 observed double mutants, more than 46% show significant negative epistasis. On the other hand, only 18 mutants (1.8%) show significant positive epistasis. Strikingly, all but one of these interactions involve a secondary mutation at position 582. As shown in Figure 2, the general pattern of epistasis is similar for most anchor data sets, with only slight differences in the curvature of the relationship between expected multiplicative fitness and observed fitness of the double mutant. Only the anchor 588F yields a clearly different pattern that more closely resembles the expected linear relationship under no epistasis. Notably, this is the only anchor that has a wild-type-like growth rate, while all other anchors have slightly deleterious fitness effects in the wild-type background. Interestingly, positive epistasis is mostly identified between slightly deleterious mutations that become wild-type-like in the presence of the anchor mutation – hence, indicating a weak compensatory effect of certain mutations at position 582. A phylogenetic alignment of 74 species across eukaryotes showed that none of the double mutations from our study were observed in the phylogeny. This is not surprising given that this region is strongly conserved, and our study contained only a very small subset of all possible double mutations. In addition, Hietpas *et al.* (2011; Figure 3, Supplementary Figure S5) studied the relationship between EMPIRIC measurements and mutational sampling in natural populations in more detail, and did not find a great correspondence. However, position 582 is the most variable position with four different amino acids observed in the phylogenetic alignment, whereas at all other positions, there are at most two different amino acids present across the phylogeny. Interestingly, in all but one of the 43 species showing a change in position 582, one or two additional changes are observed at other positions in the studied region of the protein, and all double mutations observed in the phylogenetic alignment contain a mutation at this position – providing further support for a compensatory nature of position 582.

**Figure 2:**
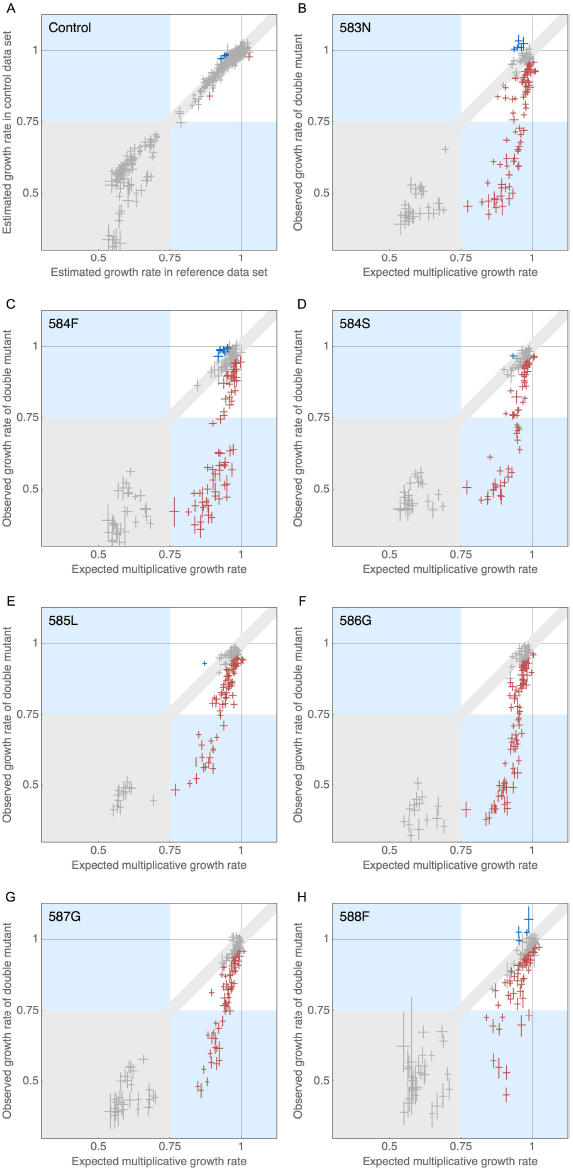
(A) Comparison of estimated growth rates between reference and control mutations that was used to determine significance level indicated by light gray area as detailed in Materials and Methods. (B)-(H) Comparison of expected and observed growth rates per anchor mutation, indicating ubiquitous negative epistasis and rare cases of positive epistasis. Data points are represented by crosses indicating the width of their respective 95% credibility intervals. Negatively epistatic interactions are shown in red, positively epistatic interactions in blue. The gray area indicates our limit of detection, as detailed in Materials and Methods.

In a comparably unbiased study focusing on an RNA recognition motif in *S. cerevisiae*, the authors found 1% positive, but only 3.6% negative epistasis in an assessment of 40,000 double mutations (Melamed et al. 2013). Interestingly, they also identified certain mutations that are prevalently involved in positive epistasis (so called “hot-spots” for epistatic interaction), while being slightly deleterious on their own. The discrepancy in the identified proportion of negatively epistatic mutations may owe to both biological and technical reasons. First, the studied RNA recognition motif may have a different underlying architecture of epistasis. Secondly, while allowing for a larger number of screened mutations, the analysis based on only two time samples in Melamed *et al.* (2013) results in higher sampling noise than our approach that is based on 5 time samples, and thus, they may have reduced resolution on epistatic effects, in particular between deleterious mutations.

### The distribution of fitness effects, from the perspective of differing genetic backgrounds

The observed pattern of epistasis is also reflected by a change in the shape of the distribution of fitness effects (DFE). We have previously reported the bimodal shape of the DFE in standard conditions (cf. Hietpas et al. 2011), which, given our categorization of strongly deleterious mutations (cf. Materials and Methods) results in a wild-type like mode harboring approximately 80% of mutations and a strongly deleterious mode harboring approximately 20% of the mutations. Figure 3 shows the DFE of each anchor data set (normalized to the anchor mutation) as compared with the single-step DFE. Although the general shape of the DFE remains bimodal, the overall shape is shifted. Firstly, the mode centered around wild-type-like fitness effects shows a smaller mean and a larger variance resulting in an increased number of deleterious mutations. Secondly, we observe a striking increase in strongly deleterious mutations, with the proportion of strongly deleterious mutants being nearly double that observed in the single-step data set. In previous work (Hietpas et al. 2013; Bank et al. 2014), we studied the change in the DFE upon altering the environment, which similarly resulted in a higher variance of the DFE, but also in an increase in beneficial mutations, consistent with expectations from Fisher’s Geometric model (FGM, Fisher 1930). In this respect, the pattern is not consistent with FGM, because the movement away from the optimum – this time achieved by anchoring a variant of lower-than-wild-type fitness – does not result in larger potential of beneficials in the DFE. We also estimated the tails shape of the beneficial tail of DFE for all data (cf. Beisel et al. 2007; Joyce et al. 2008; Bank et al. 2014) and observe a similar pattern (cf. Supplementary Figure 2), indicating a bounded tail shape for all anchor data sets, with very small estimated distances to the optimum. This suggests that the potential for adaptation is reduced when the genetic background is altered.

**Figure 3:**
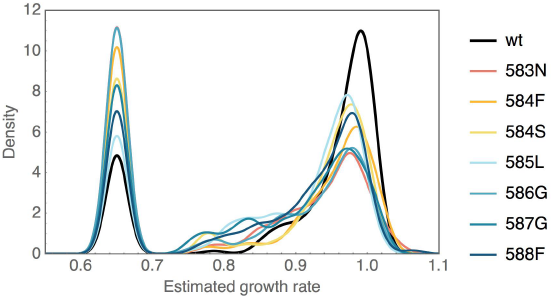
Distribution of fitness effects (DFE) of single point mutations reachable from the wild type and from each of the anchor mutations, with the most important features being an increased variance of the wild-type-like mode and a much larger proportion of strongly deleterious mutations. DFEs are plotted as smoothed histograms with a kernel width of 0.015. Strongly deleterious mutations were binned to 0.65.

The observed lower density of the DFE at the fitness of the reference type suggests that, upon a change of the genetic background away from the well-adapted wild type, there is an increased risk that a new mutation is deleterious. In the following subsection, we will argue how this observation is consistent with a concave shape of the local fitness landscape.

### The concave shape of the fitness landscape

With the above results in hand, it is feasible to consider the shape of the local fitness landscape. The curvature and ruggedness of fitness landscapes is a highly discussed topic that has recently been addressed in several ways, both empirically and theoretically (Bershtein et al. 2006; Zeldovich et al. 2007; Lunzer et al. 2010; Jiang et al. 2013; de Visser and Krug 2014).

Repeatedly, a concave relationship has been found between phenotypic effects such as stability or expression level and the resulting fitness. A concave phenotype-fitness relationship would explain two evident patterns observed in the data: Firstly, a concave fitness landscape results in the above-discussed shift in the DFE upon taking a step away from the wild type. This happens because, as the population is dislocated from the fitness plateau, fitness effects are more likely to end up in the deleterious mode of the DFE. Secondly, a concave fitness landscape results in a correlation between the expected multiplicative effect and the strength of epistasis (cf. Fig. S3), because upon dislocation of the population towards lower fitness, the slope of the fitness landscape becomes increasingly steep – resulting in a stronger discrepancy between the expected and the observed effect of a double mutation. This type of correlation has been identified and discussed in previous studies using a variety of data sets (e.g., Schenk et al. 2013; Velenich and Gore 2013).

We tested whether the observed epistasis pattern can be explained by a simple concave fitness landscape using two models that describe different relationships between an underlying phenotypic effect and the resulting fitness. Model 1 is based on a binding saturation curve as it has been used to explain the relationship between expression and fitness in Jiang *et al.* (2013),

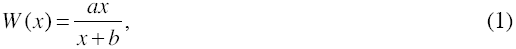

where *x* describes the phenotypic effect, *a* is the maximum possible growth rate (i.e., *a*-1 represents the distance to the optimum in Fisher’s geometric model), and *b* describes the curvature. Model 2 is based on a relationship between thermodynamic stability and fitness and was previously used to explain the epistasis pattern in Jacquier *et al.* (2013). Here,

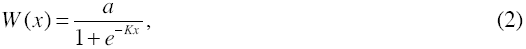

where *x* describes the phenotypic effect, *a* is the maximum possible growth rate, and *K* describes the curvature. We use the respective functions to determine an expected growth rate of a double mutant as detailed in Fig. 4, resulting in the expectations

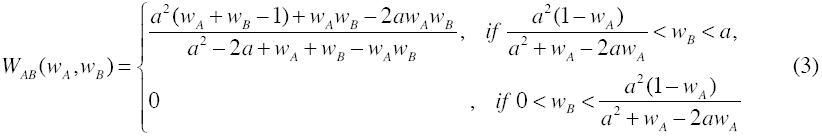

for the model based on the saturation curve (Model 1), and

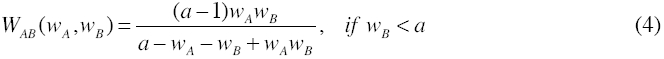

for the model based on thermodynamic stability (Model 2). Notably, both expectations are dependent on only one parameter, *a*, which is the maximum possible fitness (or, in terms of the FGM, the distance to the optimum). We fitted the data as detailed in Materials and Methods, resulting in the estimates summarized in Figure 4.

**Figure 4:**
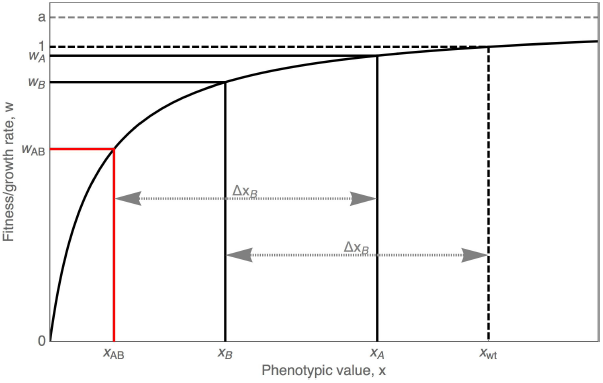
Representation of a concave relationship between phenotype and fitness. The fitting procedure is based on a reverse mapping, *X(w)*, of the growth rate of single-step mutations, *W(x)*,under the expectation that phenotypic values interact linearly, hence, *x*_*AB*_ = *x*_*A*_ − Δ*x*_*B*_ = *x*_*A*_ − (*x*_*wt*_ − *x*_*B*_) . This yields the expected growth rate of a double mutant, *w*_*AB*_, from the two measured single-step growth rates *w*_*A*_ and *w*_*B*_ (cf. also Supplementary Material Online).

Notably, both models result in errors that are orders of magnitude lower than those under the assumption of a multiplicative relationship, with the thermodynamic model resulting in slightly lower errors than the binding model (cf. Fig. 5B). However, the estimates for the distance to the optimum resulting from the binding model are more consistent with those obtained from the tail distributions (Fig. 5A; cf. Supplementary Figure 2B). In particular, the estimated maximum growth rates under the thermodynamic model are consistently lower than the highest observed growth rates for the studied data set. Therefore, while there is good support for a simple concave relationship between phenotype and fitness (under both models), it is unclear which one of the tested models provides a better explanation for the observed pattern.

**Figure 5:**
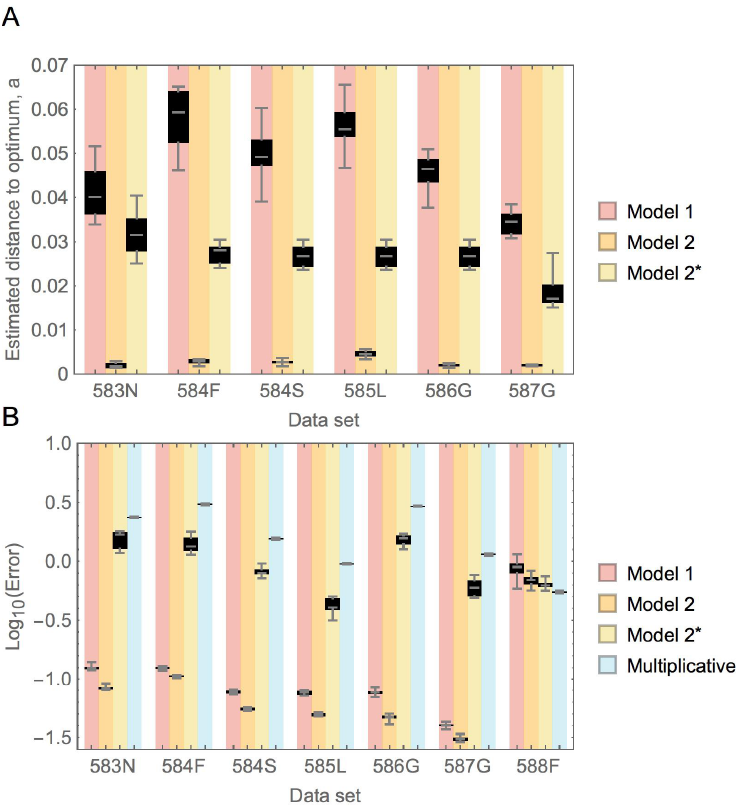
(A) Estimated parameters for the concave shape of the fitness landscape, for Model 1 based on a saturation binding curve, and Model 2 based on a thermodynamic stability curve. Because the best fit for Model 2 consistently estimated a distance to the optimum *a* (representing the maximum observable improvement over the wild type growth rate) incompatible with the mutation with the highest observed growth rate in the data set (i.e., the best observed mutation would overshoot the estimated optimum), values are added for a modified version of model 2, here termed Model 2*, that allows only estimates consistent with all observed mutations (i.e., with *a≥*max*(w_i_)-1*, where max(*w*_*i*_) denotes the maximum of all observed growth rates in the respective data set). Anchor mutation 588F is not shown, because parameter estimates diverge, supporting a multiplicative model for this mutation. Estimated distances to the optimum differ between underlying fitness curves, with the stability model yielding much higher values than the thermodynamic model. In the modified model (Model 2*), the estimated distance to the optimum is always the difference between the wild-type growth rate and the highest observed growth rate in the respective data set (i.e. *a*_*2\**_=max*(w_i_)-1)*. (B) Errors resulting from a fit to the above-described models, calculated as the minimum Euclidean distance between the estimated curve and the data. Errors are orders of magnitude smaller for the two tested models than for the multiplicative assumption, with the thermodynamic model yielding slightly lower errors than the model based on the saturation-binding curve. Only for the anchor mutation 588F that results in wild-type-like fitness, the multiplicative model yields the best result. Estimates are based on 10 samples from the posterior distributions of the estimated growth rates.

In any case, the support for a concave fitness landscape suggests that the wild-type genotype resides in a “robust” position in sequence space, in which a single mutation is less likely to result in harmful effects than if it occurred on a genetic background one mutational step away. This notion of mutational robustness has been observed and discussed previously (e.g., Denby et al. 2012; Lauring et al. 2013), with a controversial question being whether mutational robustness is a trait under natural selection (and the resulting consequences for the process of adaptation), or whether the architecture of proteins and the resulting sequence space can intrinsically confer robustness (van Nimwegen et al. 1999; de Visser et al. 2003; Draghi et al. 2010; Wagner 2012; Stern et al. 2014). While the present data set is insufficient to distinguish between these two hypotheses, future systematic studies of epistasis in different environments could be tailored to shed light on this question.

### Correlation between epistatic effects on different backgrounds

Above we have shown that there is a strong correlation between the multiplicative effect size and the amount of epistasis. This could indicate that not the two specific positions involved, but only their single effect size determine the amount of epistasis. To evaluate this hypothesis, we calculated the correlation between the same second-step mutations on different anchor backgrounds. While correlations are observed, they are stronger between classes of similar biophysical properties rather than between individual sites (see Table 3), indicating that biophysical properties rather than the net growth rate are determining the amount of epistasis. There is a strong correlation between secondary mutations with the anchors 583N, 584F, and 584S (all three at the most exposed positions), and between the anchor data sets 586G and 587G (same anchor amino acid, and both buried positions). Furthermore, we observe the weakest correlation between the anchor data set 588F and all other data sets. Notably, 588F is the data set with the highest anchor fitness, indicative of a different starting position on the fitness landscape.

### A biochemical interpretation of observed epistatic effects

Given the known structure of this region, it is possible to better interpret these results biochemically. Indeed, the anchor mutations used in this study were chosen such that 1) based on structural inspection, they were likely to alter either the exterior composition or the main-chain conformation of this region of Hsp90, and 2) they were well tolerated in the parental background (Table 1). We hypothesized that these non-conservative but well tolerated substitutions would sensitize Hsp90 to secondary mutations and thus provide insights into the interplay between mutations impacting protein conformation and/or exterior composition. We sampled anchor mutations at both solvent-accessible surface as well as solvent-inaccessible core positions. In the following, we investigate a number of hypotheses based on the biochemical properties of the tested mutations.

#### The role of core and surface positions

Core positions in proteins tend to have a dominant impact on structure and dynamics, while positions on the surface tend to play a primary role in mediating intermolecular interactions (e.g., Dill 1990; Cordes et al. 1996; Marshall and Mayo 2001). There are exceptions to this general trend because the impact of a mutation on protein stability, dynamics, and function depends on detailed atomic interactions that are not perfectly captured by surface/core classification. For example, glycine mutations typically increase protein flexibility at any position because the lack of heavy atoms in the glycine side chain provides greater access to main chain conformations than any other amino acid. To broadly span potential mutant structural effects, we chose anchor mutations (Figure 1) at two solvent exposed positions (F583N and W585L), two mutations at a glycine position (G584F and G584S), and three mutations at solvent inaccessible positions (S586G, A587G, and N588F).

In light of the solved crystal structure of Hsp90 (Ali et al. 2006), we here consider the above-described interactions in terms of the biochemical properties of this region. Based on this structure, amino acids 582-590 form a loop with two bulky hydrophobic residues projecting into solvent, as well as several other residues buried and solvent shielded in the structure. To probe the structural features of a position and their association with a particular category of epistasis, we classified each position as either solvent exposed or solvent shielded based on the proportion of solvent-exposed surface area estimated by PoPMuSiC 2.0 (Dehouck et al. 2009, reported in Table 1).

We found that combinations of surface and core mutations are more likely to exhibit negative epistasis than combinations of two surface or two core positions (cf. Figure 6A). This is consistent with previous literature proposing strong interactions between surface and core mutations in close structural proximity (Toth-Petroczy and Tawfik 2011). Previous biochemical analyses of stability indicate that solvent-shielded mutations make evolutionarily conserved energetic contributions to global folding/unfolding (Gupta et al. 2012; Das et al. 2013). Biophysical analysis of mutants within this region indicate single mutants have very little effect on global folding stability (Ohmae et al. 1998; Jiang et al. 2013). A likely explanation for the abundance of core mutations contributing to negative epistasis is that the conformation of this loop is impacted by mutations at solvent-shielded positions and this in turn impacts the positioning of surface residues, and therefore fitness.

**Figure 6:**
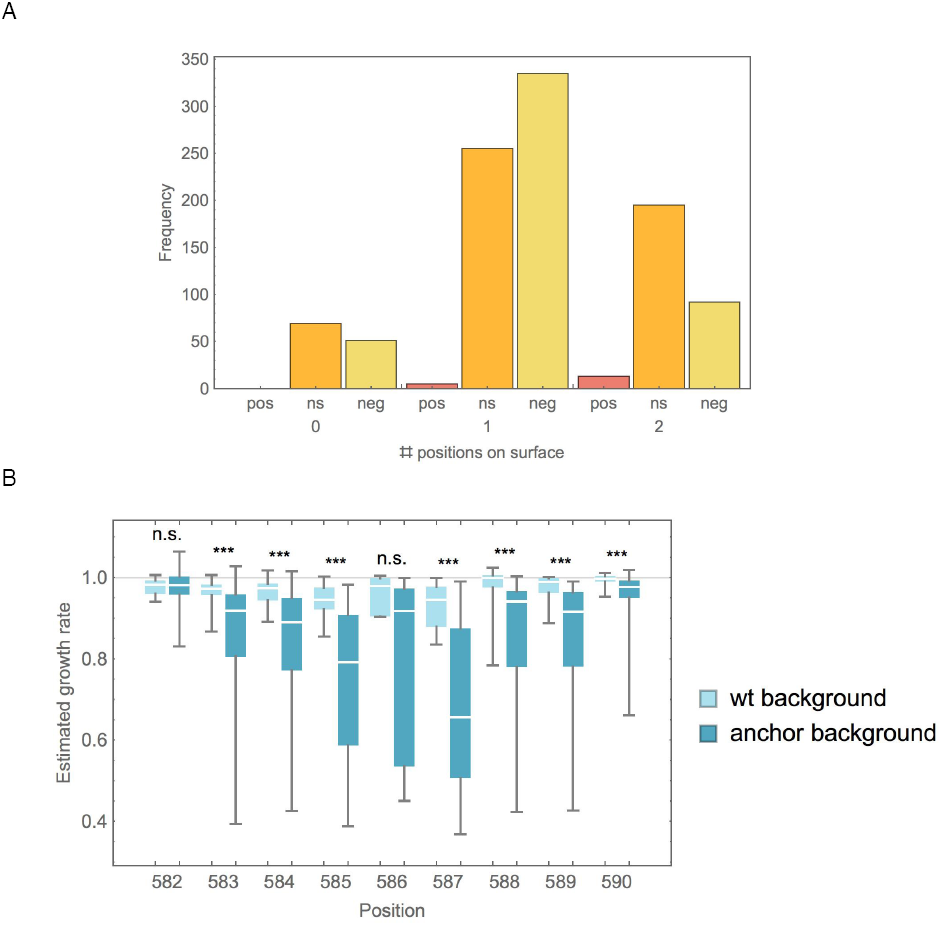
(A) Numbers of positively (pos), negatively (neg), and non-epistatic (ns) mutations, distinguished by the number of surface positions involved (strongly deleterious mutants excluded). There are significantly more negative interactions in double-mutations involving one core and one surface position than expected by chance (Binomial test, p<10^-5^), and in turn more positive (p<0.005) and non-significant (p<10^-6^) interactions between two surface positions. (B) Growth rate of mutations on wild type as compared with anchor background (mutations strongly deleterious on wild-type background are excluded). Medians are significantly different (Mann-Whitney test, p<0.001) for all but two positions. Position 582 is the prominent position exhibiting positive epistasis, whereas mutations at 586 have highly variable effects both on the wt and the anchor background.

Notably, all observations of positive epistasis included at least one solvent exposed position, and all but one instance of positive epistasis involves a mutation at position 582. This region of Hsp90 has previously been characterized as a putative protein binding interface due to characteristic hydrophobic residues projecting into solvent (Ali et al. 2006; Hietpas et al. 2011), so the overarching explanation for fitness effects within this region is based on crosstalk between the hydrophobic character of surface positions and the orientation of these residues dictated by core residue packing.

#### Position-dependent epistatic effects

Figure 6B shows the estimated growth rate of mutations on the wild-type background as compared with an anchor background for each position separately. Interestingly, the median growth rate was significantly lower on the anchor background as compared with the wild-type background for all but two positions, 582 and 586. Whereas 586 is a highly constrained position that represents a putative hydrogen-bonding site (cf. Table 1) and that results in highly variable fitness effects of different mutations, 582 is the position that is involved in all but one cases of positive epistasis identified in this study. This indicates that, while most studied positions are highly sensitive to secondary mutations, position 582 is not only less sensitive, but it can also compensate for slightly deleterious effects at other positions – an interpretation consistent with the phylogenetic observation discussed above.

## CONCLUSION

The datasets generated here provide direct measures of the fitness effects of 1,015 double amino acid substitutions including the magnitude and frequency of intragenic epistatic interactions between substitutions. The EMPIRIC approach is ideally suited to address this challenge as all mutants are introduced into the same batch of competent yeast rapidly expanded from a single colony – which critically provides for the control of the genetic background. To calculate epistatic effects, we compared direct measures of the fitness effects of double mutants to the predicted independent fitness effect of each individual mutant. We find epistatic interactions are common, and the majority of epistasis is negative in direction. Further, by focusing on neutral and weakly deleterious first step mutations, we examine an important and previously underexplored area of the fitness landscape. Unlike in previous studies that were concerned with the change in the fitness landscape caused by environmental stresses, our results are not consistent with predictions from Fisher’s Geometric model - with the prevalence and strength of beneficial mutations not necessarily increasing upon displacing the population away from the optimum. Furthermore, we quantify the changes in the DFE due to epistasis, and show that the observed pattern is consistent with a concave relationship between phenotype and fitness, indicating that the wild type resides in a position of high mutational robustness in sequence space. Finally, a biochemical interpretation of these results in light of the known structure of Hsp90 indicates a complex interplay between mutations that impact conformation and exterior facing composition.

## MATERIALS AND METHODS

### Anchored Library Generation

Seven Hsp90 point mutations (F583N, G584F, G584S, W585L, S586G, A587G, and N588F) previously observed to have wt-like to slightly deleterious fitness (growth rate within 2.5% of wt) under standard growth conditions (see Hietpas et al. 2011)were chosen as anchors. Within each of these seven anchored Hsp90 backgrounds, systematic site saturation mutagenesis was used to introduce second point mutations throughout the amino acid 582-590 region. The amino acid position fixed as the anchor was chosen to act as a single site library control to determine if the fitness effects of individual mutations were reproducible, and the preceding position maintained fixed to provide an internal measure of sequencing noise Mutagenesis was carried out as previously described (Hietpas et al. 2011).

### Yeast transformation and selection conditions

Yeast manipulations and growth competitions were performed as previously described (Jiang et al. 2013). Briefly, mutants were encoded on p417GPD, a plasmid/promoter system that closely matches the endogenous expression of Hsp90 (Nathan and Lindquist 1995). Each anchored library was separately transformed into the DBY288 strain of *S. cerevisiae* (can1-100 ade2-1 his3-11,15 leu2-3,12 trp1-1 ura3-1 hsp82::leu2 hsc82::leu2 ho::pgals-hsp82-his3).Transformed cells were amplified in medium containing galactose (SRGal -H +G418; per liter: 1.7g yeast nitrogen base without amino acids, 5g ammonium sulfate, 0.1g aspartic acid, 0.02g arginine, 0.03g valine, 0.1g glutamic acid, 0.4g serine, 0.2g threonine, 0.03g isoleucine, 0.05g phenylalanine, 0.03g tyrosine, 0.04g adenine hemisulfate, 0.02g methionine, 0.1g leucine, 0.03g lysine, 0.01g uracil 200mg G418, with 1% w/v raffinose and 1% galactose) such that wt Hsp82 protein was co-expressed along with each mutant. Transformation of the library yielded on average 110,000 individual isolates. Following the amplification of pooled transformants, cells were diluted into fresh SRGal–H +G418 medium and grown to mid log phase. Selection was initiated by transferring cells to shutoff conditions consisting of synthetic dextrose medium (SD –H +G418; identical to SRGal –H +G418 media but with 2% dextrose in place of raffinose and galactose). Yeast cells were diluted periodically (with minimum population sizes in gross excess to library diversity) to maintain log phase growth and samples isolated at different time points in shutoff conditions (Supplementary Figure S1).

### Sequencing and data analysis

Lysis, sample preparation, and sequencing were performed as previously described (Hietpas et al. 2012). In brief, plasmid DNA was harvested from yeast pellets, and the mutated region was selectively amplified by PCR and prepared with Illumina primer binding sites, and a barcode used to distinguish each time point. Sequencing was performed on the Illumina HiSeq platform at the UMass Medical School core sequencing facility. Sequencing produced 21.6million reads of 99% read confidence per base and the relative abundance of each mutant at each time point extracted as previously described (Hietpas et al. 2012).

### Data processing

Individual growth rates were estimated according to the approach described by Bank *et al.* (2014). Briefly, an outlier detection step was performed to detect sequencing errors at single data points, and subsequently, posterior probabilities were estimated using a Bayesian Monte Carlo Markov Chain (MCMC) approach. Because of our interest in the effect of amino acid mutations, nucleotides coding for the same amino acid were interpreted as replicates with equal growth rates. On the nucleotide basis, mutations resulting in HpaII sites, and mutations with a maximum read number smaller than 50 throughout all time samples were excluded. The resulting MCMC output consisted of 10,000 posterior estimates for each amino acid mutation.

### Classification of mutations

Epistasis was calculated based on 10,000 posterior samples as the difference between the observed double-step growth rate and the product of the single-step growth rates, *e*_*ab*_=*w*_*ab*_-*w*_*a*_*w*_*b*_. Significance was assigned upon determining a global mis-measurement cutoff interval I=[-0.02;0.02] resulting from an evaluation of single-step controls and comparison between single-step replicates (cf. Figure 2A) and choosing a false positive rate of 1.5%. A mutation was classified as negatively (positively) epistatic, if its 95% credibility interval was non-overlapping with the cutoff interval and the mean was smaller (larger) than 0. Because accurate measurement of very deleterious mutations is difficult and the determined growth rate dependent on the composition of the experimental library (see also Bank et al. 2014), we classified a strongly deleterious category of mutations (those with growth rate smaller that 0.75), for which we did not quantify epistasis.

### Fitting procedure

*Mathematica* 9.0.1 was used in an iterative procedure to obtain the best fit of the curves described in the main text (Equations 3 and 4). For each suggested parameter, the Euclidean distance between the data and the curve was minimized, and the resulting squared distance was used as measure of the error. Convergence of the minimization algorithm was checked based on a graphical output of the iteration procedure.

**Table 2.** Correlation of epistatic effects between anchor data sets<sup>a</sup>

| R<sup>2</sup> | 583N | 584F | 584S | 585L | 586G | 587G | 588F |
| --- | --- | --- | --- | --- | --- | --- | --- |
| 583N | 1 | 0.84 | 0.9 | 0.38 | 0.45 | 0.57 | 0.28 |
| 584F | 0.84 | 1 | 0.81 | 0.47 | 0.39 | 0.43 | 0.3 |
| 584S | 0.9 | 0.81 | 1 | 0.57 | 0.53 | 0.42 | 0.43 |
| 585L | 0.38 | 0.47 | 0.57 | 1 | 0.46 | 0.31 | 0.48 |
| 586G | 0.45 | 0.39 | 0.53 | 0.46 | 1 | 0.86 | 0.27 |
| 587G | 0.57 | 0.43 | 0.42 | 0.31 | 0.86 | 1 | 0.11 |
| 588F | 0.28 | 0.3 | 0.43 | 0.48 | 0.27 | 0.11 | 1 |

<sup>a</sup>Highest correlations are found between certain surface positions (highlighted in blue), and between glycine anchors at core positions (highlighted in red). Anchor mutation 588F with wild-type-like single-step fitness shows the lowest correlation with all

## ACKNOWLEDGMENTS

Financial support for this work was provided by grant R01-GM083038 from the National Institutes of Health to D.N.A.B., and by grants from the Swiss National Science Foundation and the European Research Council to J.D.J. C.B. is grateful to the Simons Foundation for support during a semester stay at the Simons Institute for the Theory of Computing, where parts of this work were performed, and to various participants of the semester program for helpful discussions and feedback. The computations were performed at the Vital-IT (http://www.vital-it.ch) Center for high-performance computing of the SIB Swiss Institute of Bioinformatics.

